# Ligand-induced and small molecule control of substrate loading in a hexameric helicase

**DOI:** 10.1101/069773

**Authors:** Michael R. Lawson, Kevin Dyer, James M. Berger

## Abstract

Processive, ring-shaped protein and nucleic acid protein translocases control essential biochemical processes throughout biology, and are considered high-prospect therapeutic targets. The *E. coli* Rho factor is an exemplar hexameric RNA translocase that terminates transcription in bacteria. Like many ring-shaped motor proteins, Rho activity is modulated by a variety of poorly understood mechanisms, including small molecule therapeutics, protein-protein interactions, and the sequence of its translocation substrate. Here, we establish the mechanism of action of two Rho effectors, the antibiotic bicyclomycin and nucleic acids that bind to Rho’s ‘primary’ mRNA recruitment site. Using SAXS and a novel reporter assay to monitor the ability of Rho to switch between open-ring (RNA loading) and closed-ring (RNA translocation) states, bicyclomycin is found to be a direct antagonist of ring closure. Reciprocally, the binding of nucleic acids to its N-terminal RNA recruitment domains is shown to promote the formation of a closed-ring Rho state, with increasing primary site occupancy providing additive stimulatory effects. This study establishes bicyclomycin as a conformational inhibitor of Rho ring dynamics, highlighting the utility of developing assays that read out protein conformation as a prospective screening tool for ring-ATPase inhibitors. Our findings further show that the RNA sequence specificity used for guiding Rho-dependent termination derives in part from an intrinsic ability of the motor to couple the recognition of pyrimidine patterns in nascent transcripts to RNA loading and activity.

**SIGNIFICANCE:** Many processive, ring-ATPase motor proteins rely on substrate-dependent conformational changes to assist with the loading of client substrates into the central pore of the enzyme and subsequent translocation. Using the *E. coli* Rho transcription terminator as a model hexameric helicase, we show that two distinct ligands – the antibiotic bicyclomycin and pyrimidine-rich nucleic acids – alternatively repress or promote, respectively, the transition of Rho from an open, RNA-loading configuration to a closed-ring, active helicase. Our findings explain several mechanisms by which Rho activity is controlled, and provide a general illustration of how intrinsic and extrinsic factors can regulate ring-type ATPase dynamics through diverse mechanisms.

## INTRODUCTION

Ring-shaped hexameric helicases and translocases are motor proteins that control myriad essential viral and cellular processes. Many hexameric motors undergo substrate-dependent conformational changes that couple activity to the productive binding of client substrates (1-4). Internal regulatory domains and exogenous proteins or small molecules frequently impact client substrate recruitment and engagement by these enzymes (5-8); however, it is generally unclear how such factors control helicase or translocase dynamics.

Rho is a hexameric helicase responsible for controlling approximately 20% of all transcription termination events in bacteria (9). Rho is initially recruited to nascent transcripts in an open, lockwasher-shaped configuration (Figure 1A) (10, 11), where it binds preferentially to pyrimidine-rich sequences (termed <u>R</u>ho <u>u</u>tilization of <u>t</u>ermination sequences, or ‘*rut*’ sites) using a ‘primary’ RNA-binding site located in the N-terminal OB folds of the hexamer (12-14). Following *rut* recognition, Rho converts into a closed-ring form (Figure 1B), locking the RNA strand into a ‘secondary’ RNA binding site (formed by two conserved sequence elements known as the Q and R loops (15)) located within the central pore of the hexamer. This conformational change, which we show in an accompanying paper to be both RNA- and ATP-dependent (16), rearranges residues in the Rho ATP binding pockets into a hydrolysis-competent state (17). Once engaged, Rho maintains primary site contacts with the *rut* sequence as it translocates 5′ to 3′ along the RNA strand in an ATP-dependent manner, a process known as tethered tracking (18-21). Rho elicits termination by applying direct or indirect forces to RNA polymerase (22, 23), dislodging it from DNA and the newly made RNA in the transcription bubble.

**Figure 1.**
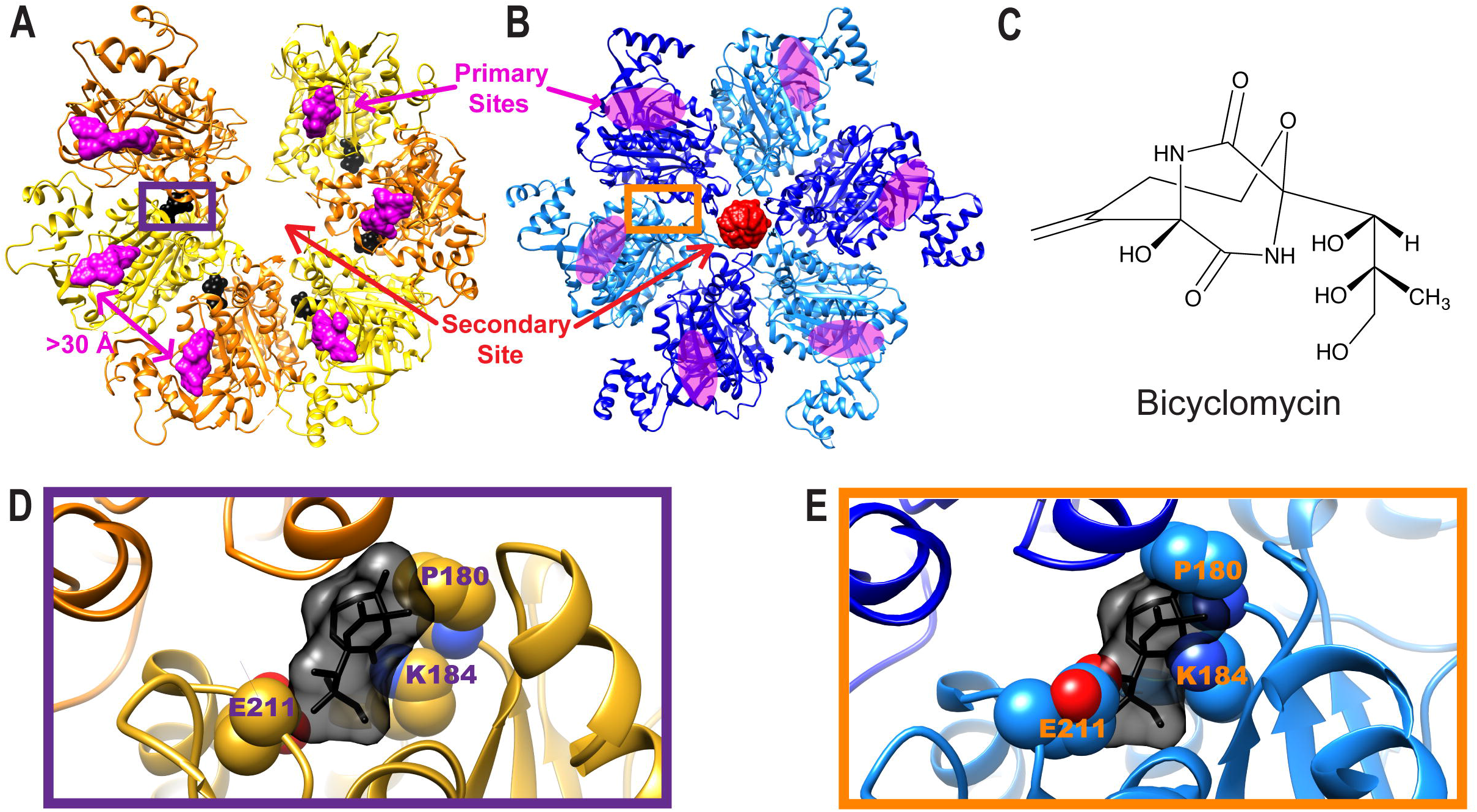
Bicyclomycin is an inhibitor of the Rho helicase. **(A)** Crystal structure of open-ring and bicyclomycin-bound Rho (PDB ID: 1XPO (24)). Rho subunits are alternatingly colored yellow and gold for distinction, bicyclomycin is black, and primary site RNAs are magenta. **(B)** Crystal structure of closed-ring and translocation-competent Rho (PDB ID: 3ICE (17)). Rho subunits are alternatingly colored cyan and dark blue, and RNA bound in the secondary site is red. Magenta-colored ovals denote the location of the primary sites that are not occupied in this crystal form. **(C)** Chemical structure of bicyclomycin. **(D)** Close-up view of the bicyclomycin-binding pocket shows that bicyclomycin nestles into a small pocket between Rho subunit interfaces. **(E)** Modeling of bicyclomycin into the closed-ring Rho structure shows clear steric clashes between the drug and Rho.

Although the basic steps of Rho-dependent transcription termination have been largely elucidated, and it has become increasingly clear that Rho’s activity, like that of many helicases and translocases, can be controlled by a variety of intrinsic and extrinsic factors. The small molecule bicyclomycin (Figure 1C), a highly specific chemical inhibitor of Rho that has been shown to wedge itself in between subunits of an open Rho ring (Figure 1D and **Supplementary Figure 1**) (24), is one such example. Previous studies have shown that bicyclomycin is a noncompetitive inhibitor of Rho ATPase activity and a mixed inhibitor of RNA binding to Rho’s secondary site (25, 26). Although subsequent structural studies suggested that bicyclomycin antagonizes Rho by sterically preventing the binding of the nucleophilic water molecule that initiates ATP hydrolysis (24), how bicyclomycin might interact with catalytically-competent Rho states, such as those thought to accompany ATPase activity and translocation (17), has not been defined.

It is similarly unclear how other dissociable factors – e.g., regulatory proteins and/or nucleic acids – influence Rho activity. It is well established that the sequence of the RNA itself has a pronounced impact on whether a transcript will be acted upon by Rho (12, 27-29). Binding of pyrimidine-rich sequences to the N-terminal RNA binding domains of Rho is a particularly well-known accelerant of Rho’s ATPase activity (30), with pre-steady-state ATPase assays showing that the formation of a catalytically competent Rho ring is governed by a rate-limiting RNA- and ATP-dependent conformational change (8). The ligand-dependence of this isomerization event correlates with the requirements for ring closure identified in an accompanying study (16). These findings raise the intriguing possibility that the sequence specificity of Rho-dependent termination is in part due to an increased efficiency of ring closure when the N-terminal RNA binding domains are occupied by pyrimidine-rich sequences.

Here we show that bicyclomycin and primary-site occupancy affect the structural state of Rho in an opposing manner. Using small-angle X-ray scattering and a fluorescence-based assay to track ring status *in vitro*, we demonstrate that bicyclomicin inhibits RNA binding to the central pore of Rho by sterically impeding ring closure. In contrast, the binding of pyrimidine-rich nucleic acids to the Rho N-terminal RNA binding domains promotes Rho ring closure, helping the capture of non-ideal, purine-rich RNA sequences within Rho’s translocation pore. Collectively, these findings help highlight diverse mechanisms by which ligand binding to discrete sites on ring-type ATPases can activate or repress conformational changes and govern the function of the motor.

## RESULTS

### The bicyclomycin-binding pocket collapses upon Rho ring closure

During the course of examining the bicyclomycin-binding pocket across different Rho structural states ((17) and (16)), we noticed that the site appeared substantially smaller in closed-ring states than in the open-ring states. Computational analysis confirmed that the pocket volume is substantially larger in the open-ring conformation (272 Å^3^, **Supplementary Figure 2A**) than in the closed-ring state (56 Å^3^, **Supplementary Figure 2B**), a trend that held true with all open- and closed-ring Rho structures that have been observed to date (**Supplementary Table 1** and **Supplementary Figure 2**). Correspondingly, alignment of a bicyclomycin-bound open ring structure (PDB ID 1XPO (24)) with a closed-ring Rho model (PDB ID 3ICE (17)) revealed extensive steric clashes with bicyclomycin and residues P180, K184, and E211 in the closed-ring state that would appear to preclude binding (compare Figure 1D and Figure 1E). From these observations, we postulated that bicyclomycin might not sterically block the attacking water needed for ATP hydrolysis, as previously proposed, but instead antagonize the RNA- and ATP-dependent conformational switching of Rho from an open-to closed-ring state.

### Bicyclomycin inhibits Rho ring closure over a range of nucleotide concentrations

To determine the impact of bicyclomycin on Rho conformational state, we first employed Small-Angle X-Ray Scattering (SAXS) to directly monitor structural dynamics in solution. An accompanying study demonstrates that open-ring Rho molecules close in the presence of RNA and the nonhydrolyzable ATP analog ADP•BeF_3_, but not in the presence of either ligand individually (16). Since bicyclomycin is a relatively modest inhibitor of Rho ATPase activity (I_50_ = 20 μM (25)), we reasoned that its effects on ring closure might be most evident when examined over a range of nucleotide concentrations. As a control, we carried out SAXS studies using a series of ADP•BeF_3_ concentrations in the presence of RNA and the absence of bicyclomycin. By plotting the average intensity of the observed scattering (I) as a function of scattering vector (q) for a variety of ADP•BeF_3_ concentrations (Figure 2A), nucleotide-dependent differences in curve shapes were clearly evident, similar to those reported in the accompanying study ((16) and **Supplementary Figure 3**). Visual inspection of the curves over a q range from 0.07 to 0.13 Å^-1^(the region of the curve in which nucleotide-dependent differences are most pronounced, Figure 2A, *Inset*) revealed that the profiles converged between 1.5 and 4 mM ADP•BeF_3_, suggesting that under these conditions Rho exists predominantly as a closed ring. Indeed, quantification of the percentage of open *vs*. closed states using a Minimal Ensemble Search (‘MES’, (31)) indicated that 1.5 mM ADP•BeF_3_ was sufficient to close the majority of Rho rings in solution (78% closed), with an increase in the percentage of closed rings observed at 2, 3 and 4 mM ADP•BeF_3_ (89%, 93% and 95% closed respectively; see Figure 2C and **Supplementary Table 2**). Similar trends were also observed in the reduction of the radius of gyration (‘R_g_’) at increasing ADP•BeF_3_ concentrations, consistent with the expected nucleotide-dependent compaction of the Rho ring upon transition to a predominantly closed-ring state (**Supplementary Figure 4**, **Supplementary Figure 5** and **Supplementary Table 3**).

**Figure 2.**
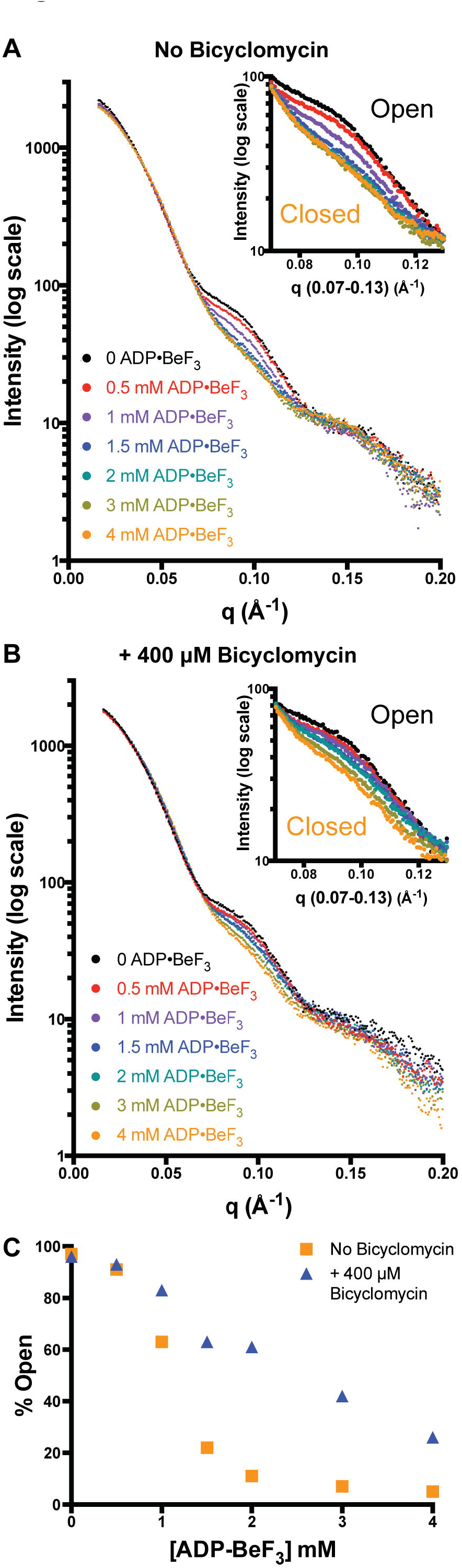
Bicyclomycin inhibits Rho ring closure over a range of nucleotide concentrations. **(A)** Scaled SAXS intensities of Rho in the presence of one rU_12_ RNA per hexamer and a variety of concentrations of the ATP mimetic, ADP•BeF_3._ **(Inset)** Zoom-in of the mid-q region, in which differences between the various SAXS curves aremost pronounced. **(B)** Scaled SAXS intensities of Rho pre-incubated with 400 μM bicyclomycin, and then combined with one rU_12_ RNA per hexamer and a variety of concentrations of ADP•BeF_3._ **(Inset)** Zoom-in of the mid-q region reveals a skew of the SAXS curves towards the open state in the presence of bicyclomycin. **(C)** Quantification of the curves represented in **(A)** and **(B)** by MES shows a clear skew towards the open ring state in the presence of bicyclomycin.

After establishing a SAXS regime for looking at changes in ring state, we next set out to determine whether bicyclomycin would antagonize the ability of the Rho ring to close in a nucleotide-dependent fashion. We first pre-incubated Rho with a high concentration of bicyclomycin (400 μM final concentration, an 8-fold molar excess over Rho monomer), and then added RNA and a variety of ADP•BeF_3_ concentrations. The resulting SAXS curves (Figure 2B) also displayed differences that were clearly dependent upon nucleotide concentration; however, at intermediate concentrations of ADP•BeF_3_ (0.5, 1 and 1.5 mM), curves were skewed towards the open-ring state relative to the drug-free curves (compare Figure 2B, *Inset* and Figure 2A, *Inset*). Quantification of ring state by MES showed a substantial shift toward the open form when bicyclomycin was present, which was particularly notable at 1.5 and 2 mM ADP•BeF_3_ (Figure 2C and **Supplementary Table 2**). This bicyclomycin-dependent inhibition of Rho ring closure was also consistent with observed increases in R_g_ upon addition of the drug (**Supplementary Figure 4A**, **Supplementary Figure 6** and **Supplementary Table 3**).

### Rho ring state can be controlled by varying bicyclomycin concentration

After observing that bicyclomycin inhibits ring closure, we wondered whether it would be possible to control the Rho ring state by varying the concentration of bicyclomycin with Rho prior to adding RNA and ADP•BeF_3._ To address this question, we elected to conduct a SAXS experiment in the presence of 1.5 mM ADP•BeF_3_, a nucleotide concentration that based on our previous titration is sufficient to close a substantial fraction (but not all) of the Rho rings present in solution. Pre-incubating Rho with varying concentrations of bicyclomycin yielded significant differences in the observed SAXS curves (Figure 3A), and by focusing on the mid-q region (Figure 3A, *Inset*), it is immediately evident that the ring state can be affected by bicyclomycin. Quantification of the percentage of the open *vs*. closed ring populations by MES (Figure 3B and **Supplementary Table 4**) shows that ring state changed from nearly completely closed (<10% open) in the absence of bicyclomycin to almost exclusively open at high bicyclomycin concentrations (79%, 85% and 90% open and 800, 1600 and 3200 μM bicyclomycin, respectively). Observed R_g_ values ranged from 46.3 Å in the absence of bicyclomycin to 48.2 Å in the presence of 3200 μM bicyclomycin, consistent with bicyclomycin concentration-dependent inhibition of Rho ring closure (**Supplementary Figure 4B**, **Supplementary Figure 7** and **Supplementary Table 5**).

**Figure 3.**
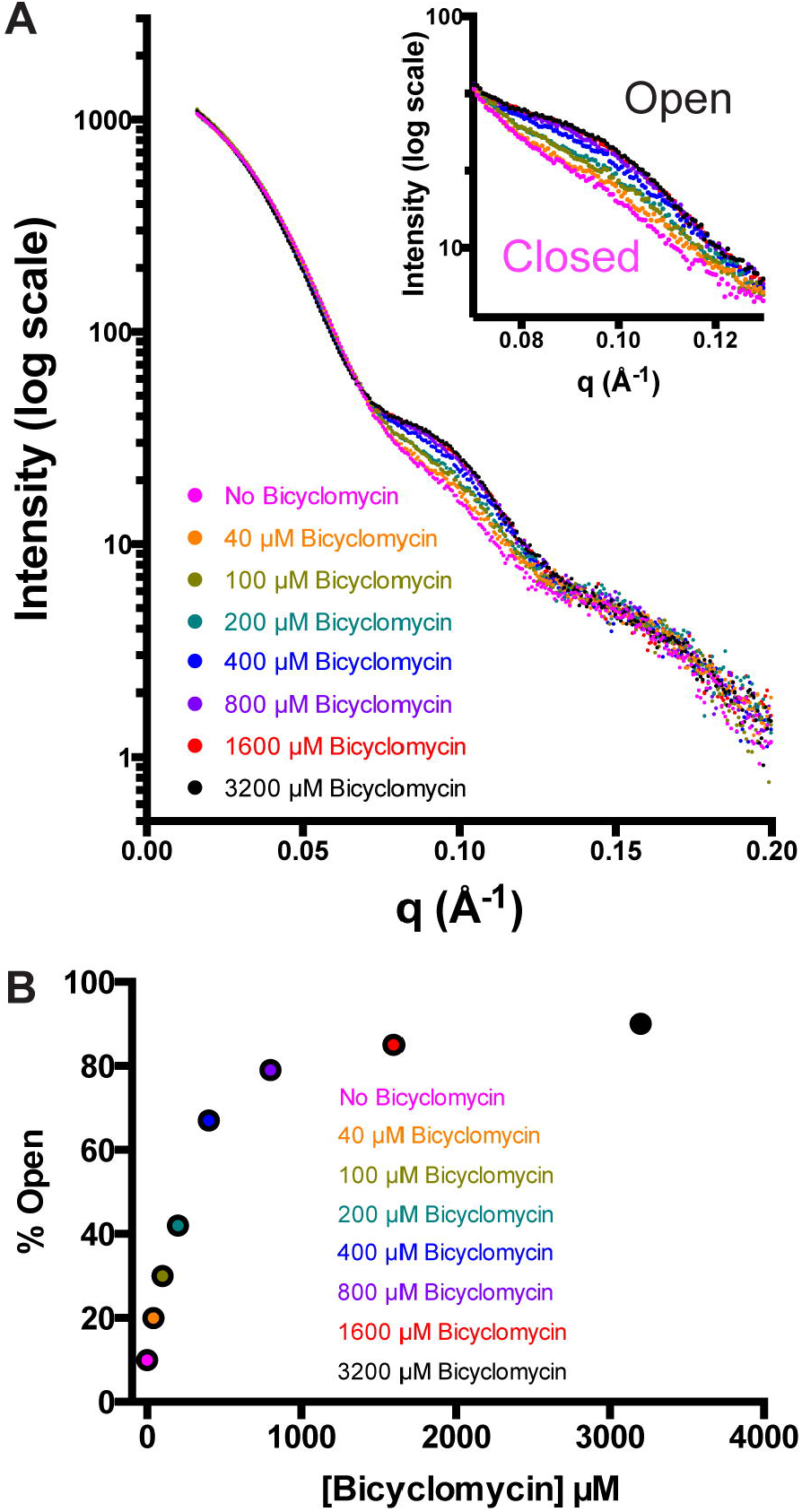
Rho ring state can be controlled by varying bicyclomycin concentration. **(A)**Scaled SAXS intensities of a variety of bicyclomycin concentrations, prior to adding one rU_12_ RNA per Rho hexamer and 1.5 mM ADP•BeF_3._ **(Inset)** Zoom-in of the mid-q region shows that ring state can be titrated from closed to open by increasing bicyclomycin concentration. **(B)** Quantification of the curves in **(A)** shows a clear titration of Rho ring state that is dependent upon bicyclomycin concentration.

### A fluorescence-based assay tracks ligand-dependent effects on Rho ring status in vitro

As SAXS studies constitute only one assessment of Rho state, we next set out to develop an *in vitro* assay to further characterize ligand-dependent effects on the conformation of the hexamer. We reasoned that a short, fluorescein-derivatized RNA would have a significantly faster tumbling rate when free in solution than when stably bound to the secondary (translocation pore) site inside a closed Rho ring (see diagram in Figure 4A), thus enabling us to track ring closure by monitoring changes in Fluorescence Anisotropy (‘FA’). To rule out the possibility that labeled RNA binding to Rho’s primary (RNA-recruitment) site might convolute the signal, we first pre-incubated the protein with a short pyrimidine-rich DNA (‘dC_5_’) at a concentration equimolar to that of Rho monomer (polypyrimidine DNAs bind both tightly and selectively to the primary sites of Rho (12, 30)). No anisotropy changes were observed with a fluorescein-labeled poly-U RNA (‘rU_12_*’) in the absence of nucleotide and in the presence of a primary site competitor (Figure 4B, black), indicating that rU_12_* alone does not have high affinity for the secondary site in open-ring Rho. By contrast, robust rU_12_* binding was observed as nucleotide was added (Figure 4B), showing that the secondary site becomes capable of stably binding RNA as the Rho ring closes. These findings are consistent with the SAXS studies carried out in the accompanying study (16), showing that both RNA and nucleotide are needed to cooperatively promote ring closure in Rho.

**Figure 4.**
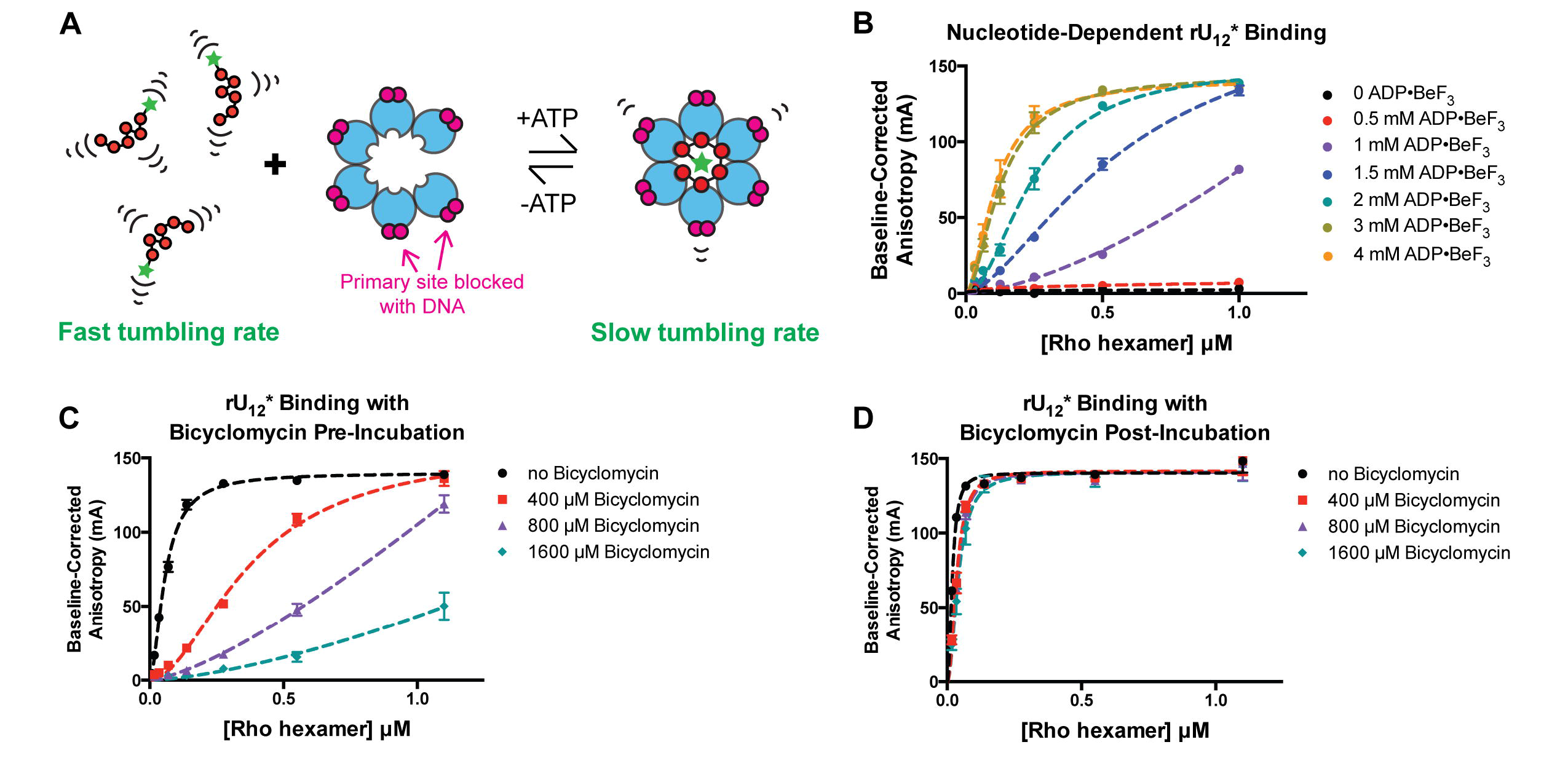
A Fluorescence Anisotropy-based RNA binding assay to track Rho ring closure *in vitro*. **(A)** Schematic illustrating how the tumbling rate of a short fluorescein-tagged RNA is slowed by capture within the Rho ring. The Rho primary sites were blocked with short DNAs, represented in magenta. **(B)** Binding of rU_12_* to Rho is dependent upon ADP•BeF_3_ concentration. **(C)** Pre-incubation of Rho with bicyclomycin before addition of RNA and ADP•BeF_3_ demonstrates bicyclomycin concentration-dependent inhibition of rU_12_*binding. **(D)** Incubation of bicyclomycin with a pre-closed Rho ring has little effect on rU_12_* binding.

To further characterize the mechanism by which bicyclomycin inhibits Rho, we next conducted an order-of-addition experiment examining the impact of bicyclomycin on rU_12_* binding either before or after Rho ring closure. As expected, pre-incubation of bicyclomycin with Rho before the addition of RNA and nucleotide inhibited rU_12_* binding in a bicyclomycin concentration-dependent manner (Figure 4C). Conversely, the impact of bicyclomycin on rU_12_* binding was far less evident when closed rings were pre-assembled before addition of the drug (Figure 4D). These data indicate that although bicyclomycin is able to bind to an open Rho ring and prevent ring closure, the molecule is unable to force open a pre-closed Rho ring.

### Rho ring closure is promoted by the binding of pyrimidine-containing nucleic acids to the N-terminal, primary site domains

Although RNA binding to Rho’s secondary site (the helicase translocation pore) is a major effector of ring closure, RNA binding to Rho’s primary site (its OB folds) also has been implicated in controlling Rho function (27). To directly probe the role of primary site ligands in modulating ring closure, we tested whether the presence or absence of dC_5_ could impact the binding of rU_12_* to the secondary site. We also conducted the same experiments with fluorescein-labeled poly-A RNA (‘rA_12_*’), which is a weaker secondary site ligand than poly-U and is incompatible with primary site binding (14, 32). A clear increase for rA_12_* affinity was evident in the presence of dC_5_ (**Supplementary Figure 8**), especially at low ADP•BeF_3_ concentrations. Under all nucleotide concentrations tested, rU_12_* bound more tightly to Rho than rA_12_* (**Supplementary Figure 8**), with dC_5_ subtly promoting ring closure at moderate (1 mM ADP•BeF_3_) nucleotide concentrations. Primary site occupancy appeared to have either no effect (1.5 mM ADP•BeF_3_) or a slight inhibitory effect (2-3 mM ADP•BeF_3_) on rU_12_* binding at higher nucleotide concentrations. Collectively, these results signify that nucleic acids bound to the primary site promote ring closure under conditions that are sub-optimal for RNA binding to the secondary site.

Based on the spacing of the OB folds in the Rho structure and the observation that the isolated N-terminal domain only engages a pyrimidine dinucleotide (Figure 1a-b) (10, 14), each dC_5_ substrate would be expected to bind one primary site at a time. Having observed that this short oligonucleotide can nonetheless promote Rho ring closure, we wondered whether longer DNAs capable of bridging multiple primary sites in a single Rho hexamer would have a more pronounced impact on ring state (see diagram in Figure 5A). We repeated the rU_12_* and rA_12_* binding assays with either a 15-mer DNA capable of bridging two primary sites (dC_15_) or a 79-mer DNA sufficiently long to potentially bridge all six primary sites (dC_75_), using an ADP•BeF_3_ concentration (1 mM) at which primary-site binding of DNA has the most significant effect on RNA binding to the secondary site. The number of putative sites that could be occupied by the DNAs was held constant relative to the dC_5_ experiments by adjusting the relative molar ratios between Rho and the DNA (**Methods**). For both rU_12_* and rA_12_*, inclusion of dC_15_ or dC_75_ more robustly promoted RNA binding than the shorter dC_5_ oligo (Figure 5B-C). Using fluorescein-labeled derivatives of the various primary-site ligands (denoted as dC_5_ *, dC_15_*, and dC_75_*), titration experiments revealed that DNAs long enough to bridge multiple primary sites have a much higher affinity for Rho than dC_5_ (Figure 5D). By fixing Rho at a concentration insufficient to drive ring closure with rU_12_* or rA_12_* and varying the concentration of primary site ligands, we found that ring closure can be driven by sufficiently high concentrations of dC_5_, dC_15_ or dC_75_ (Figure 5E-F). These data indicate that primary site occupancy favors binding of both optimal (poly-pyrimidine) and non-optimal (poly-purine) RNAs to Rho’s secondary site.

**Figure 5.**
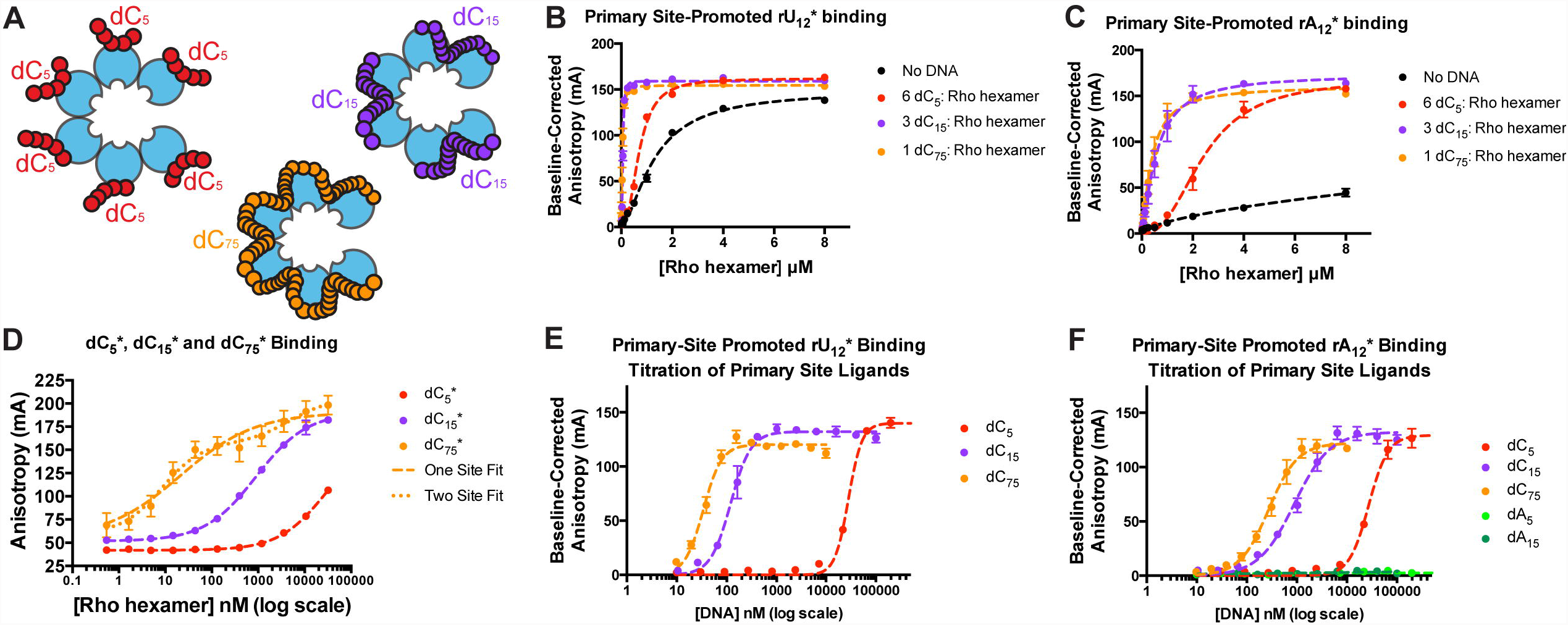
Primary site occupancy promotes Rho ring closure. **(A)** Cartoon representation of how dC_5_, dC_15_ and dC_75_ could occupy one, two, or six primary sites, respectively. **(B)** Binding of rU_12_* to Rho is promoted by polycytidylic DNAs of various lengths. **(C)** Binding of rA_12_* to Rho is promoted by polycytidylic DNAs of various lengths. **(D)** Longer DNAs (dC_15_* and dC_75_ *) capable of bridging multiple primary sites progressively promote secondary site binding more effectively than does a short DNA (dC_5_ *) that can only occupy one primary site. **(E)** Binding of rU_12_* can be fully driven at saturating concentrations of dC_5_, dC_15_ or dC_75._ **(F)** Binding of rA_12_* can be fully driven at saturating concentrations of dC_5_, dC_15_ or dC_75_. Polypurine DNAs (dA_5_ and dA_15_, light and dark green, respectively), which do not bind Rho’s primary site, do not facilitate Rho ring closure.

## DISCUSSION

Ring-type helicases and translocases control numerous essential biological processes (33, 34). Although generally capable of standalone motor activity, multiple lines of evidence are increasingly highlighting complex mechanisms by which these enzymes are regulated. Protein-protein and protein-ligand interactions, small molecule effectors, and post-translational modifications have all been implicated in controlling disparate types of ring-ATPases; how a majority of these factors exert their various stimulatory or antagonistic effects is generally not well understood.

In the present study, we discovered that bicyclomycin, one of the few small molecule agents known to inhibit a ring-type motor, acts on its target – the Rho transcription termination factor – by sterically blocking a conformational change from an open-to a closed-ring state. Rho ring closure, which an accompanying study shows is dependent upon both ATP and secondary-site bound RNA (16), triggers RNA strand engagement and rearranges residues in the ATP binding pocket into a hydrolysis-competent state. By comparing a bicyclomycin-bound, open-ring conformation of Rho to other structures of Rho sub-states, we found that the drug-binding pocket collapses upon ring closure (Figure 1). Follow-up SAXS studies show that bicyclomycin does not simply occlude binding of a catalytic water as previously proposed (24), but instead directly impedes nucleotide-dependent closure of the Rho ring (Figures 2-3). Using a FA-based assay to track ring closure *in vitro*, we further observed that bicyclomycin acts on pre-opened Rho rings, and has little effect after ring closure (Figure 4). Since inhibition of secondary-site RNA binding is indicative of bicyclomycin binding to Rho (26), these results strongly indicate that bicyclomycin is unable to bind to pre-closed Rho rings. Thus, bicyclomycin appears to inhibit Rho by stabilizing a conformation that is incapable of ATP hydrolysis and incompatible with stable RNA binding to its motor regions.

In the course of investigating bicyclomycin’s effects on Rho, we also found that binding of nucleic acids to Rho’s primary, mRNA recruitment site promotes the switching of Rho from an open-to closed-ring state. This finding is in accord with prior studies showing that Rho’s ATPase activity can be stimulated by primary site occupancy (30), and that Rho’s ATPase rate is in part regulated by an RNA- and ATP-dependent conformational change (8), presumably into a closed-ring state (16, 17). The inclusion of polypyrimidine DNAs, which bind selectively to the primary site and also promote ATPase activity (30), increased Rho’s affinity for both rU_12_ and rA_12_ substrates at nucleotide concentrations that are otherwise too low to stably drive ring closure (**Supplementary Figure 8**). This result suggests that the binding of a *rut* sequence to Rho’s primary site, which is formed by an N-terminal OB-fold in the protein (12, 13), allosterically promotes ring closure.

We further found that DNAs capable of bridging two (dC_15_) or six (dC_75_) primary sites within the Rho hexamer promoted ring closure more robustly than short (dC_5_) DNAs capable of only occupying one primary site (Figure 5A-C). The more robust impact of these longer DNAs is likely due to higher affinity for Rho (Figure 5D) rather than by a particular signal that might be propagated by a nucleic acid segment bridging multiple primary sites, as all three ligands drive complete ring closure at sufficiently high concentrations (Figure 5E-F). We also found that ring closure was more sensitive to primary site occupancy with rA_12_ than with rU_12_, which is of interest because polypurines bind to the secondary site more weakly than polypyrimidines (18). Collectively, these results suggest that the RNA recruitment sites on Rho work together to count the number and spacing of pyrimidine recruitment motifs in an mRNA, and promote ring closure around non-optimal secondary site binding sequences such as purine-rich regions (Figure 6). Future studies looking at different types of di-pyrimidine patterns on both natural and synthetic substrates will be needed to probe how this interrogation occurs at a molecular level.

**Figure 6.**
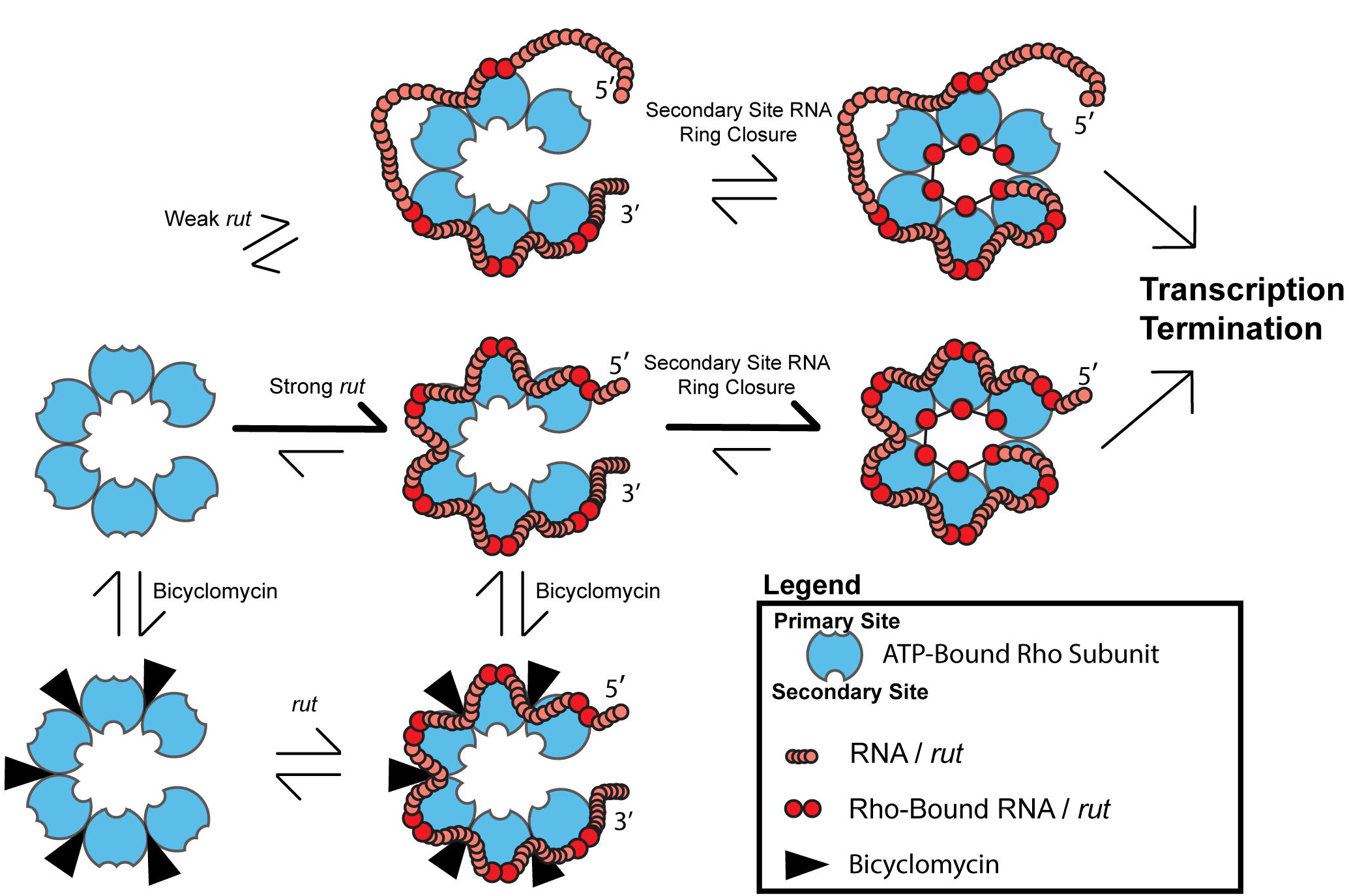
Model for how bicyclomycin and primary site ligands regulate Rho ring closure, and thereby the capacity of the helicase to promote transcription termination. After a *rut*-containing mRNA first binds to the Rho primary sites and subsequently the secondary site, the Rho ring snaps shut to trap the mRNA within the central pore. This conformational change to a translocation-competent closed ring state is less efficient in the presence of a weakly binding *rut* sequence with non-optimal di-pyrimidine number/spacing (top), and is completely blocked by bicyclomycin (bottom, black wedges). Open-ring Rho must bind to the *rut* sequence via primary site interactions and close before its ATPase activity can promote 5′ to 3′ RNA translocation and transcription termination.

In using an auxiliary ligand-binding element to aid activity, Rho joins a growing list of processive ring helicases and translocases that are subject to both intrinsic and extrinsic regulation. These factors include the bacterial DnaB helicase (6), the MCM2-7 helicase (7, 35), and the proteasomal Rpt1-6 ATPases (5). In each of these motors, a built-in accessory domain or a dissociable set of adaptor proteins is used to choreograph proper substrate recruitment and engagement with motor activity. How control linkages between the accessory elements and the motor regions are established through long-range binding and allosteric transitions remains a frontier question.

In closing, it is worth noting that there is substantial interest in identifying small molecule approaches to inhibiting ring-ATPases (36). Unfortunately, because ATP binding pockets are often well conserved across different motor families, developing such inhibitors has been challenging. Interestingly, Rho is one of the few ring-type motor proteins for which a small molecule with therapeutic properties has been identified. Here we have shown that this agent, bicyclomycin, does not antagonize the chemistry of the ATPase reaction, but instead acts as a conformational inhibitor of ring dynamics, entering a binding pocket that is only accessible during a portion of the Rho mRNA loading and translocation cycle. These findings suggest that by using appropriate screens for ring dynamics, coupled with counter screens against general ATPase activity, it may be possible to identify new generations of selective inhibitors with clinical potential.

## METHODS

### Calculation of bicyclomycin-binding pocket volumes

Bicyclomycin-binding pocket volumes in various Rho structures were calculated using the program POVME, using a spherical search with an 8 Å radius and default parameters (37). The center of the bicyclomycin binding pocket was defined as the location of the C1 atom of bicyclomycin in Chain B of the structure 1XPO (24), and the corresponding site in all other structures as determined through superposition to this site. Waters were removed from all structures before calculation of binding pocket size.

### Small-angle X-ray scattering (SAXS) sample preparation

Rho was purified as described previously (17), concentrated to 60 mg ml^-1^, and then dialyzed overnight into SAXS collection buffer (150 mM KCl, 50 mM HEPES-NaOH (pH 7.5), 5 mM MgCl_2_, 5% Glycerol). Aliquots of bicyclomycin (obtained as a generous gift from Dr. Y. Itoh and the Fujisawa Pharmaceutical Co., Ltd., Japan) were generated through the addition of dialysis buffer to the lyophilized drug. Rho samples were then diluted with bicyclomycin aliquots (for drug containing samples) or dialysis buffer (for drug-free samples) and incubated together overnight. RNA (obtained as 250 nmol syntheses of rU_12_ from IDT) was initially suspended in DMPC-treated RNase-free water (MP Biomedicals), supplemented with an equal volume of 2× SAXS buffer (300 mM KCl, 100 mM HEPES-NaOH (pH 7.5), 10 mM MgCl_2_, 10% Glycerol) immediately before use, and added at a concentration equimolar to that of Rho hexamer. ADP•BeF_3_ aliquots were generated using a 1:3:15 ratio of ADP:BeCl_2_:NaF and diluted with an equal volume of 2× SAXS buffer. Final SAXS samples were distributed into an Axygen 96-well PCR plates, and obtained through the sequential addition of 8 μl of Rho/bicyclomycin, 8 μl of RNA, and finally 8 μl of ADP•BeF_3_ after a two-hour wait. Plates were then flash frozen in liquid nitrogen and stored at −80°C.

### Small-angle X-ray scattering (SAXS) data collection and analysis

Frozen plates were thawed in water, spun for 1 min at 1000 × g, and data collected at 12°C using an automated, high-throughput system at BL 12.3.1 at the Advanced Light Source (38). Buffer-subtracted curves were generated using an automated Ogre script provided by ALS BL12.3.1 (Greg Hura). We focused our analysis on data from 1 s exposures of 2.5 mg ml^-1^ Rho samples because the buffer-subtracted curves showed the least evidence of radiation damage or aggregation under these conditions. Data scaling, Guinier analyses and P(r) calculations were conducted using the ScÅtter software suite (39). MES calculations were conducted using the FOXS web server (40), using default parameters with the open (PDB ID 1PV4 (10)) and closed (PDB ID 5JJI (16)) ring Rho structures provided as inputs. Unbuilt regions of the open and closed ring structures were modeled in using alignments with Rho structures in which these regions were visualized. All plots were generated using the Prism 6 software suite.

### Flourescence Anisotropy (FA)-based RNA binding data collection and analysis

Rho was concentrated to 60 mg ml^-1^ and dialyzed overnight into SAXS buffer. DNA oligonucleotides (dC_5_, dC_15_ and “dC_75_” (dC_15_TC_15_TC_15_TC_15_TC_15_)) utilized as primary site ligands were purchased from IDT, were resuspended in the SAXS buffer used for dialysis, and concentrations were calculated based on A_260_ readings and extinction coefficients provided by IDT. For assays, Rho was diluted with SAXS buffer in the presence or absence of DNA, and left to incubate for >1 h on ice. 5′ 6-Fluorescein amidite-labeled rU_12_ and rA_12_ oligonucleotides were purchased from IDT, resuspended in MilliQ-purified water to 100 μM and stored as small aliquots at −80° C until used. Rho was pre-incubated with fluorescein-labeled oligonucleotides, BSA and DTT (20 nM, 0.5 mg ml^-1^ and 5 mM final concentrations respectively) in SAXS buffer on ice for at >30 min, and then combined with ADP•BeF_3_ in SAXS buffer for >30 min before reading (referred to herein as assembly of closed rings). In the case of pre/post bicyclomycin incubations, Rho was either incubated with bicyclomycin for 1 h before assembly of closed rings (“pre-incubation”), or Rho was incubated with bicyclomycin for >2h after assembly of closed rings (“post-incubation”). All FA measurements were made at 30° C using a Biotek Neo2 plate reader in Corning 384-well low volume plates (#3544). Baseline corrections (where appropriate) and averaging of duplicates into single points were calculated using a custom Python script; graphs were created in the Prism 6 software suite, and fits generated by nonlinear regression to one-site binding models (with the exception of dC_75_*, which fit notably better to a two-site model – here, both one- and two-site fits are shown on the corresponding plot).

## ACKNOWLEDGEMENTS

The authors thank members of the Berger lab and Andreas Martin’s lab (UCB/HHMI) for helpful discussions and careful reading of the manuscript. The authors also thank Greg Hura, Michal Hammel, Nathan Thomsen and ALS Beamline 12.3.1 for guidance on SAXS sample preparation and data processing. This research was supported by funding from the NIGMS (GM017474 and GM066698), the G. Harold and Leila Y. Mathers Foundation (to JMB) and a NSF Graduate Research Fellowship (to MRL).

## AUTHOR CONTRIBUTIONS

MRL and JMB designed experiments. MRL and KD collected SAXS data. MRL analyzed SAXS data, and both collected and analyzed RNA binding data. MRL and JMB wrote the paper.

